# Visual contributions to the perception of speech in noise

**DOI:** 10.1101/2025.09.26.678636

**Authors:** L.C. Alampounti, H. Cooper, S. Rosen, J.K. Bizley

**Affiliations:** Ear Institute, University College London, London, United Kingdom; Speech, Hearing and Phonetic Sciences, University College London, London, United Kingdom; Audiology Department, Royal Berkshire NHS Foundation Trust, Reading, United Kingdom

**Keywords:** audiovisual integration, auditory scene analysis, temporal processing, speech perception in noise, lipreading, audiovisual benefit, coordinate response measure, CRM

## Abstract

Investigations of the role of audiovisual integration in speech-in-noise perception have largely focused on the benefits provided by lipreading cues. Nonetheless, audiovisual temporal coherence can offer a complementary advantage in auditory selective attention tasks. We developed an audiovisual speech-in-noise test to assess the benefit of visually conveyed phonetic information and visual contributions to auditory streaming. The test was a video version of the Children’s Coordinate Response Measure with a noun as the second keyword (vCCRMn). The vCCRMn allowed us to measure speech reception thresholds in the presence of two competing talkers under three visual conditions: a full naturalistic video (AV), a video which was interrupted during the target word presentation (Inter), thus, providing no lipreading cues, and a static image of a talker with audio only (A). In each case, the video/image could display either the target talker, or one of the two competing maskers. We assessed speech reception thresholds in each visual condition in 37 young (≤ 35 years old) normal-hearing participants. Lipreading ability was independently assessed with the Test of Adult Speechreading (TAS). Results showed that both target-coherent AV and Inter visual conditions offer participants a listening benefit over the static image audio-only condition, with the full AV target-coherent condition providing the most benefit. Lipreading ability correlated with the audiovisual benefit shown in the full AV target-coherent condition, but not the benefit in the Inter target-coherent condition. Together our results are consistent with visual information providing independent benefits to listening, through lip reading and enhanced auditory streaming.

## Introduction

Listening in noise is difficult, so much so that our ability to cope with interactions in noisy settings has interested researchers since Cherry (1953) defined the ‘cocktail party problem’. It is known that listening is not only supported by the auditory system, but also by vision. Sumby and Pollack, (1954) in their seminal work, demonstrated that looking at the talker’s face can augment participants’ speech perception when speech was being perceived in the presence of background noise. More recent work has shown that that visual information lowers the signal-to-noise ratio (SNR) for which speech can be perceived at a fixed level of performance (Bernstein et al., 2004; Grant & Seitz, 2000), as well as increase the percentage of correct stimulus identification at fixed SNRs (Tye-Murray et al., 2007) when compared to the performance in audio-only conditions. Other advantages of audiovisual speech perception (compared to audio only), include better comprehension of stories (Arnold & Hill, 2001), more efficient shadowing of spoken passages (Reisberg et al., 1987), reduced cognitive effort to perform the auditory task at hand (Anderson Gosselin & Gagné, 2011), and improved ability to learn degraded speech (Wayne & Johnsrude, 2012). The “visual enhancement” of audition described in these studies is referred to as the audiovisual benefit (AV benefit).

Studies of speech-in-noise perception largely focus on contributions of lipreading, that is, the visually-conveyed phonetic information obtained from the talker’s mouth movements. Growing evidence, however, demonstrates that audiovisual temporal coherence alone – that is, temporal coherence between (both naturalistic, and artificial) auditory stimuli and artificial visual stimuli – can also offer listening benefits (Atilgan & Bizley, 2021; Maddox et al., 2015; Yuan et al., 2020, 2021; but see Capelloni et al., 2023). Such benefits can be obtained with auditory stimuli coupled to disks of fluctuating size, thought to mimic the natural temporal correlations between the amplitude envelope of vocalisations and mouth opening size (Chandrasekaran et al., 2009) and is indicative of a bottom up, language-independent process (Bizley et al., 2016). That such benefits are observed for non-speech stimuli, and when visual stimuli are themselves uninformative is evidence that they arise through enhancing auditory selective attention, either through augmenting the ability to segregate competing sounds, or to correctly select the correct sound in a mixture. It remains unclear whether listeners simultaneously exploit both audiovisual temporal coherence and lipreading during speech perception, largely due to difficulties isolating audiovisual temporal coherence using conventional speech-in-noise tests (Lee et al., 2019). Even if both mechanisms contribute simultaneously to AV benefit, their relative contributions remain to be determined.

Thus, the aims of this study were to two-fold. Firstly, to develop a speech-in-noise test that is capable of measuring individuals’ AV benefit provided by looking at a talker’s mouth when listening in noise and estimate the magnitude of lipreading as opposed to other contributions. Secondly, to test the hypothesis that visual benefits arise through both lipreading and enhancing auditory scene analysis mechanisms.

## Methods and Materials

### Development of the vCCRMn speech-in-noise test

To capture an AV benefit, a speech-in-noise test must include a condition where both audio and visual (through video) cues are available to listeners, providing a contrast with an audio-only condition. Moreover, to measure contributions from both lipreading and audiovisual temporal coherence mechanisms, the speech-in-noise test must be designed so that the two mechanisms do not confound each other. While lipreading contributions are independent of the presence of sound, audiovisual temporal coherence requires sound (Bizley et al., 2016; Lee et al., 2019). Consequently, the test must be designed such that participant performance does not saturate through the use of vision alone, but, instead, allows for the effective integration of sound cues as well. At the same time, since visual benefits to listening are known to be most beneficial during adverse listening conditions (Helfer & Freyman, 2005) the listening task must be sufficiently hard such that visual cues do not become redundant.

The audio-only Children’s Coordinate Response Measure speech-in-noise test (CCRM; Messaoud-Galusi et al., (2011); see also (Bolia et al., (2000) and Brungart et al., (2001) for the Coordinate Response Measure, or CRM, corpus from which the CCRM was inspired) served as the basis for developing our speech-in-noise test. The CCRM was selected as a starting point due to its widespread use in research with both adults and children, as well as its clinical applicability (Bianco et al., 2021; Billings et al., 2024; Messaoud-Galusi et al., 2011; Saleh et al., 2023).

The CCRM sentences follow the template “Show the [animal] where the [colour] [number] is,” where an animal word is chosen from [dog, cat, cow, duck, pig, sheep], the colour from [black, blue, green, pink, red, white], and the number from [1, 2, 3, 4, 5, 6, 8, 9]. Thus, all animal, colour and number words are monosyllabic. The target sentences always use the animal “dog” (i.e., “Show the dog where the [colour] [number] is”), while the remaining sentences, which feature other animals, colours, and numbers from those of the target and from each other, are used as maskers. The participants’ task is to identify the colour and number words mentioned in the target sentence.

The CCRM was adapted to include a video component, making the vCCRM. Initial pilot results revealed that the default choice of monosyllabic colours and numbers rendered the task too easy for some participants, as those target words could be identified accurately via lipreading alone. Consequently, in the new version of the vCCRM, the second key word was changed from a number to a monosyllabic noun. In designing the final set of 16 nouns, the mouth movements for the onset and offset of each noun were considered, ensuring that no noun exhibited unique mouth movements. In this way, each noun was visually confusable with at least two other nouns in the set – sharing a common beginning with one and a common ending with another. Pilot results showed that the task could not be resolved through silent lipreading alone; rather, both lipreading and auditory cues were necessary for successful performance, making the task a promising tool for the study.

This adapted version is hereby referred to as the *video version of the CCRM with nouns* (or *vCCRMn*). Similarly to the CCRM, sentences followed the template “Show the [animal] where the [colour] [noun] is”. Animal options included [cat, cow, dog, duck, pig, sheep], colour options included [black, blue, green, pink, red, white] and noun options included [bed, bin, boat, bone, bowl, cloak, clock, coat, cone, lock, oak, oat, pear, peg, pen, pin].

Each trial of the vCCRMn was comprised of three simultaneously presented sentences – one target and two maskers – featuring different animal, colour, and noun combinations each. The target sentence always included the animal “dog.” Trials featured either a male or female target talker, with maskers that included one male, and one female talker, which were different from the target and from each other. The talkers presented were randomly drawn from a pool of four possible talkers (two male and two female). A total of 2304 sentence stimuli were recorded at UCL’s anechoic chamber – 576 for each of the two male and two female talkers, all native speakers of Southern British English. All recorded speakers gave permission for the use of their image in the test, and in any resulting publications or presentations. Continuous audio was sampled at 44100 Hz and segmented into 3 second segments, with the speech aligned to the end of the segment. Sentences were variable in duration depending on the talker, and aligning at the end ensured that the sentence onset times were variable across competing talkers, and the colour and noun words of the target were always masked by the non-target talkers.

The task included three main conditions. The naturalistic condition ‘Audiovisual’ (AV) presented participants with both the video and the audio of a talker and was expected to provide AV benefits that encompass the contributions of both lipreading and audiovisual temporal coherence (see Figure 1A for an example trial of the AV condition, and Figure 1C for an example of the response panel). The ‘Interrupted’ (Inter) condition differed from the AV condition in that the entire screen froze during the [colour] [noun] combination, thereby removing lipreading cues during that period. Consequently, the Inter condition was expected to eliminate any contributions to performance of directly lip reading the target words, but leave intact the cues that allowed the observed talker to be tracked through the mixture. Finally, the task included an ‘Audio-only condition with a static image’ (A), which served as a performance baseline for assessing listening benefits. Each of the three main conditions was further divided into two sub-conditions. In the target-coherent sub-condition, the voice of the target talker matched the face presented in the video, whereas, in the masker-coherent sub-condition, the video presented matched one of the two masker voices (Figure 1B). This was also true for the static image audio-only condition, thus, also allowing us to assess potential advantages of looking at the static image of the target talker as compared to one of the two masker talkers.

**Figure 1:**
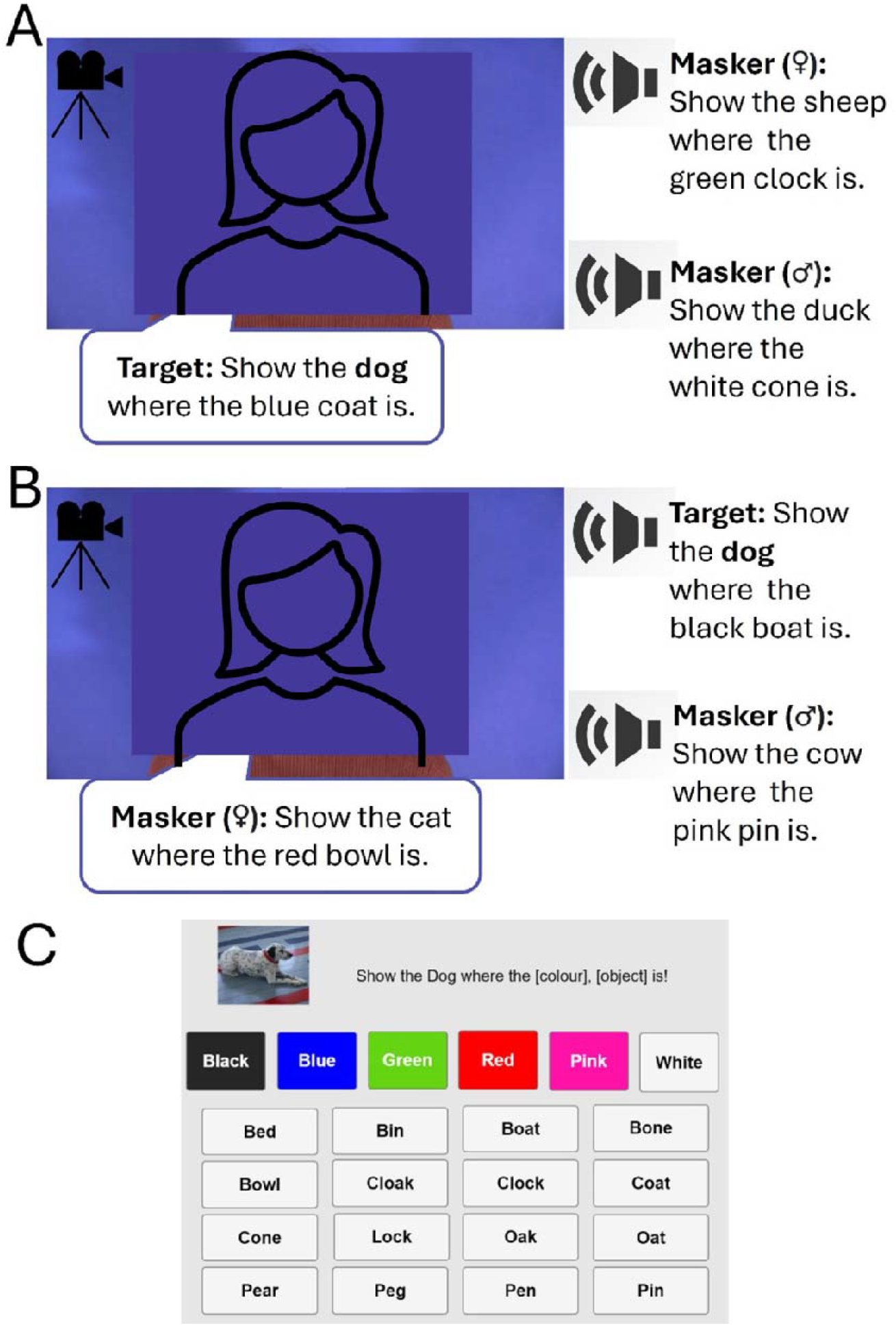
[note: figure edited to remove photos for bioRxiv submission]. The video version of the Children’s Coordinate Response Measure task, using nouns (vCCRMn). A) Example of an Audiovisual (AV) target-coherent trial. Participants were presented with the video of the talker whose voice was made the target talker voice in that trial (i.e., the talker uttering the sentence including the animal “dog”). Their task was to follow the target talker’s voice and identify the colour and noun key words in it. Two masker voices uttering a sentence of the same form but featuring different animal, colour, and noun words were presented in the background. All four recorded talkers (two male, two female) could be either target or masker talkers across conditions. B) Example of an AV masker-coherent condition. The video presented matches the voice of one of the masker speakers, while participants are still tasked with identifying the key words of the target sentence. C) Response panel with colour and confusable nouns set. The Interrupted video (Inter) and Static Image (A) conditions followed the same structure.

### Participants

Participants were recruited via the UCL Ear Institute and the UCL SONA subject pool. The experiment was conducted under the protocol approved by the University College London Research Ethics Committee (ref. 3866/003). Informed written consent was obtained from all participants prior to their participation. They were all compensated for their time.

A total of 37 participants (ages 19-35; Mean = 25.92, SD = 4.17), all native speakers of British English, were recruited and tested in person. The sample size was determined via a power calculation that accounted for potentially small effects of audiovisual temporal coherence on the AV benefit (Cappelloni et al., 2023; Maddox et al., 2015). Based on preliminary observations, an effect size of 1 dB SNR (difference between the target-coherent A and target-coherent Inter conditions) and a standard deviation of 2 dB SNR were assumed. With a Type I error probability of 0.05 and a power of 0.80, the minimum required sample size was estimated to be 33.

Inclusion criteria required that participants were native speakers of British English, had normal hearing status (see Pure-tone audiometry), normal or corrected-to-normal vision (including normal colour vision), the ability to maintain focus for the duration of the study, absence of neurological, psychiatric, or developmental conditions, no prior training in lipreading and no intrusive tinnitus, no language-related difficulties, no exposure to loud sounds within the 24 hours preceding participation and no participation in the previous pilot versions of the vCCRMn.

### Equipment and apparatus

All testing procedures were completed inside a triple-walled Industrial Acoustics Company (IAC) soundproof booth located at the Human Labs of University College London’s Ear Institute. The speech-in-noise and lipreading tests were run on a Dell latitude laptop. Prior to testing, adjustments were made using an adjustable desk, chair and laptop stand so that the screen was comfortably positioned at an eye level. Participants wore closed-back, around-the-ear headphones (Sennheiser, HD 280 Pro Dynamic HiFi Stereo), which were tested and calibrated using an artificial ear (Bruel and Kjaer Type 4153) and a measuring amplifier (Bruel and Kjaer, 3110-003) to ensure a sound level output of 65 dB SPL (RMS normalised). Participants were instructed to maintain their eye-gaze on the task display during task testing. A camera positioned on top of the screen – connected to a computer in the experimenter’s control room – was used to monitor compliance with this instruction. Communication between participants and the experimenter was maintained at all times via a microphone installed within the IAC booth.

Participants’ hearing status was assessed with a calibrated audiometer (Interacoustics, AS629, with Radioear DD450 headphones).

### Test battery

Participants carried out an array of tests during their testing session. The primary performance measure was the audiovisual speech-in-noise task *vCCRMn*, as detailed below. Additional measures included a Pure-tone audiometry (PTA) test and a silent lipreading test (TAS; Mohammed et al., (2006)). All measures were completed within a single < 2-hour testing session.

### vCCRMn

The vCCRMn was implemented as described above (‘Development of the vCCRMn speech-in-noise test) and used an adaptive one-up one-down (1U1D) staircase procedure. This tracked participants’ speech reception threshold-fifty (SRT_50_) – i.e. the SNR at which a participant is expected to complete 50% of the trials successfully. The staircase maintained the same overall level across trials and adjusted both the level of target and masker sentences to adjust the SNR. Beginning with an SNR of 20 dB, staircase step sizes were 8 dB SNR until the first track reversal, then 6 dB SNR until the second, 4 dB SNR until the third, and 2 dB SNR for all subsequent reversals. The sequence was set to stop after four runs of the final step size had been concluded or when a total of thirty trials had been completed. Participant SRT_50_s for each run of a vCCRMn condition were estimated as the mean of the last four staircase reversals.

Participants completed three runs of each of the three main conditions of the task (AV, Inter, and A), with the conditions presented in a random order across runs. Within each run two simultaneous staircases operated, one with target-coherent trials and one for masker-coherent trials, which alternated randomly and were not cued to the participant. Each run took approximately 5 mins and yielded two threshold estimates (one for target-coherent and one for masker-coherent visual trials). Overall participant performance for each coherence type of each condition was estimated as the mean of the three thresholds (one per run) obtained for that participant. Communication between participants and the experimenter was maintained at all times via a microphone installed within the IAC booth.

Before embarking on testing, participants were given a short demonstration that included example trials from each of the six vCCRMn sub-conditions. Following the demonstration, participants completed a five-minute practice session consisting of 8 trials. Each trial presented a randomly-selected vCCRMn condition and a randomly-selected vCCRMn sentence at 20 dB SNR, spoken by any one of the four talkers, and could be either target- or masker-coherent. Participants passed the practice session if they correctly identified the target words in at least 7 out of the 8 trials. They were given two opportunities to pass the practice session. Not passing the practice session led to the end of their participation and compensation for their time.

### Pure-tone audiometry

Each participant’s hearing was assessed via air conduction PTA, following the British Society of Audiology’s guidelines (British Society of Audiology, 2018). Briefly, hearing thresholds were measured at the frequencies 1000, 2000, 4000, 8000, 500, and 250 Hz in the order listed. Hearing status was, then, determined as the mean of the better-ear scores (measured frequencies between 500 and 4000 Hz). Participants with a PTA threshold of ≤ 20 dB HL were classified as having normal hearing (see also Figure 2).

**Figure 2:**
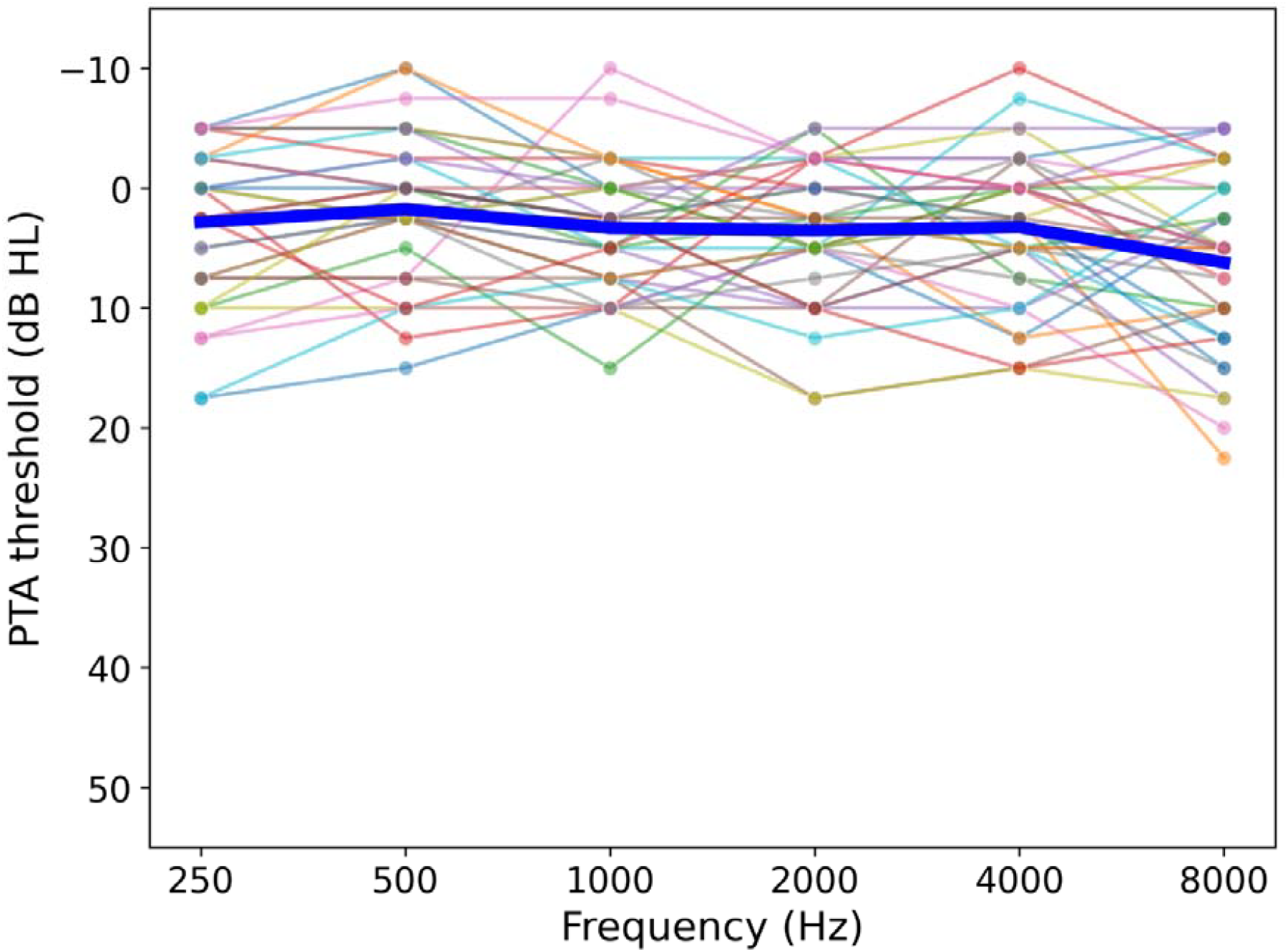
Mean PTA threshold (averaged across both ears) as a function of frequency. Each line represents a participant. The thick central line depicts the mean performance across all participants.

### Silent lipreading task

Silent lipreading ability was independently measured with the Test for Adult Speechreading (TAS). Detailed information on the test – including instructions on how to conduct the test – is available in Mohammed et al., (2006) and on the TAS webpage (https://dcalportal.org/tests/tas). Briefly, the test consists of video clips of silent speech produced by either a male or a female speaker. It includes three core tests of speechreading ability (assessing words, sentences and short stories), and three additional subtests (minimal pairs, sentence stress, and question or statement). Participants completed all core and subtest tasks. Lipreading scores (percentage correct) on the sentence-level test were selected to represent participants’ lipreading ability, as this test revealed neither floor nor ceiling effects.

### Data Analysis

Visual benefits were computed from the SRT_50_s for the AV and Inter conditions relative to the A condition. Specifically, the AV benefit was estimated using the formula:

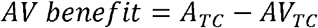

Where TC indicates target-coherence. Similarly, the Inter benefit was computed as:

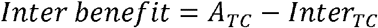

Coherence-type benefits were computed from the differences between masker- and target-coherent condition pairs. For the AV, the coherence benefit was estimated as:

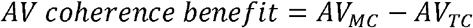

Where MC indicates masker-coherence, and TC indicates target-coherence (as defined above). Similarly, the Inter coherence benefit was computed as:

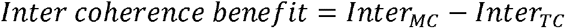

Finally, the static image-coherence benefit was computed as:

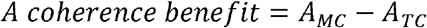

A 3 × 2 repeated measures ANOVA was used to examine the effects of visual condition (AV, Inter, and A), visual coherence type (Target, Masker), and their interaction on participant performance in the vCCRMn task. Post-hoc comparisons were conducted using pairwise paired t-tests with Bonferroni correction. Cohen’s d and mean differences between contrasted conditions were computed measures of effect size.

Pearson correlations were performed to assess the relationship between AV benefit and Inter benefit, AV benefit and lipreading score, and Inter benefit and lipreading score.

All statistical analyses were conducted in R (Version 4.2.0).

## Results

We developed a version of the CCRM that required that listeners report the colour and noun spoken by a target talker in the presence of two maskers, in three audiovisual conditions: full video (AV), interrupted video (Inter), and static image with audio only (A).

### Listeners benefit from visual information

Participants thresholds varied across visual stimulus conditions (Figure 3). A 3 x 2 repeated measures ANOVA yielded significant main effects for both visual condition (F(2, 210) = 7.28, p < 0.001) and visual coherence type (F(1, 210) = 29.89, p < 0.001), as well as a significant interaction between the two (F(2, 210) = 18.27, p < 0.001). To further explore this interaction, we conducted post-hoc paired t-tests.

**Figure 3:**
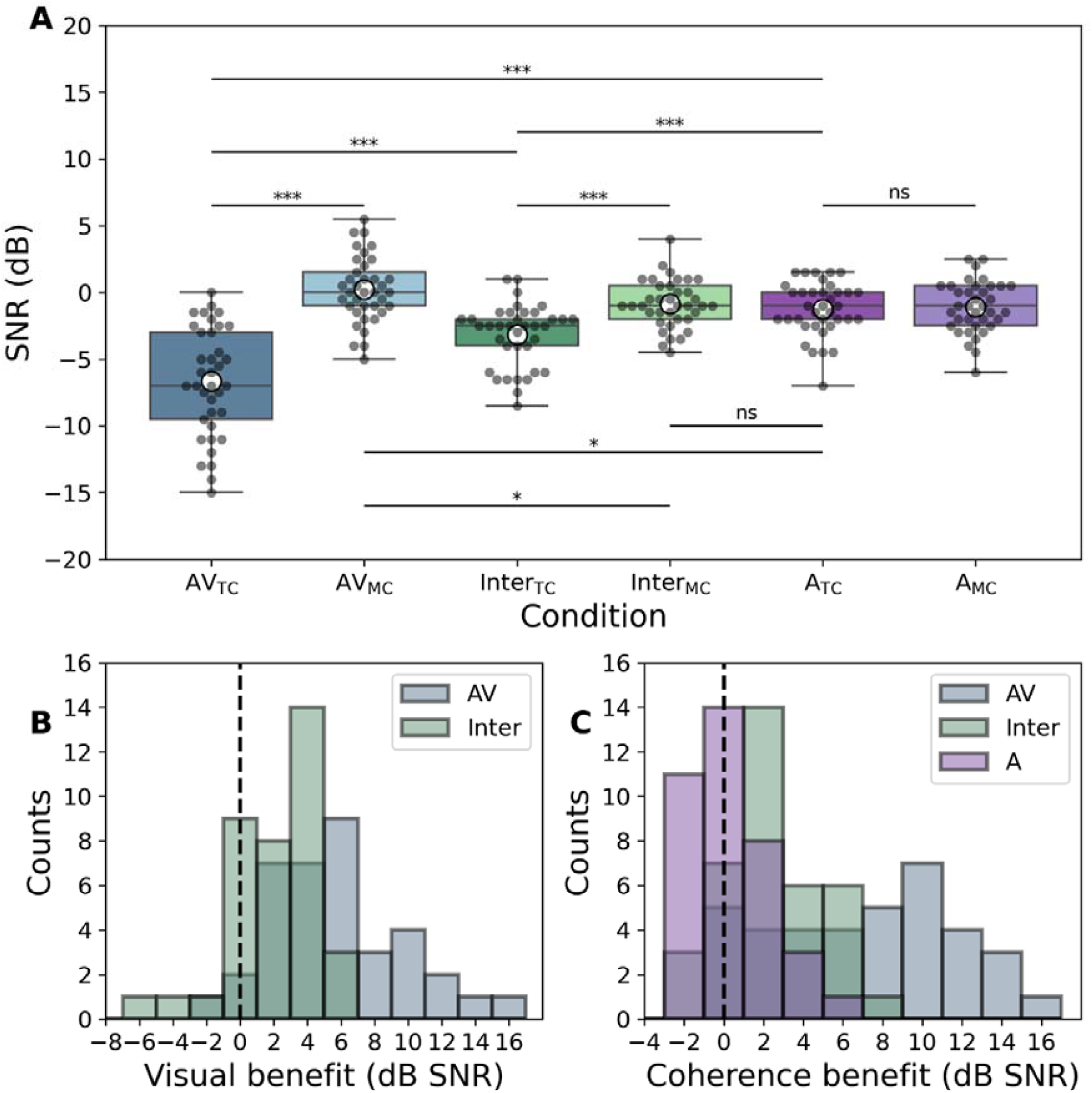
The effect of visual condition on speech-in-noise performance and assessment of benefits. A) Boxplots displaying performance in each of the six vCCRMn sub-conditions. White dots depict mean performance, boxes indicate the quartiles, whiskers the distribution range, and grey dots are individual participants. Pairwise comparisons are shown as horizontal lines over the relevant conditions (* indicates p < 0.05; *** indicates p < 0.001; ns = non-significant). TC = target-coherent condition; MC = masker-coherent condition. B) Histograms of visual benefits for the AV and Inter conditions. C) Histograms of coherence type benefits for the AV, Inter, and A conditions.

These analyses showed that performance was significantly better in the AV_TC_ condition compared to both the A_TC_ baseline (AV benefit = 5.4 dB SNR) and the Inter_TC_ (difference = 3.5 dB SNR); performance in the Inter_TC_ condition was also significantly better than in the A_TC_ (Inter benefit = 1.9 dB SNR) (see also Figure 3A). For both AV_TC_ and Inter_TC_, thresholds were significantly lower than the respective masker-coherent conditions (AV coherence benefit = 6.9 dB SNR; Inter coherence benefit = 2.3 dB SNR). Coherence type did not influence performance in the static image (A) conditions. Finally, performance was significantly worse in the AV masker-coherent condition than the audio only (AV detriment = 1.45 dB SNR). Full details, including t-test statistics, p-values and effect sizes for all comparisons are shown in Table 1.

**Table 1:** Summary of the post-hoc paired t-test comparisons. The comparisons are categorised by type. Measures of effect size, including Cohen’s d and mean differences between the conditions being compared are also included. Star symbols in the p-value column denote statistical significance (* indicates p < 0.05; *** indicates p < 0.001). TC = target-coherent condition; MC = masker-coherent condition.

| Comparison type | Condition 1 | Condition 2 | t-statistic | DoF | Bonferroni corrected p-value | Cohen's d | Mean difference (dB SNR) (Condition 1 – Condition 2) |
| --- | --- | --- | --- | --- | --- | --- | --- |
| Visual condition comparisons | AV | Inter | -2.66 | 36 | 0.028 * | -0.31 | -1.2 |
|  | AV | A | -3.76 | 36 | < 0.001 *** | -0.44 | -2.1 |
|  | Inter | A | -2.81 | 36 | 0.019 * | -0.33 | -0.8 |
| Visual benefits | AV<sub>TC</sub> | A<sub>TC</sub> | -8.16 | 36 | < 0.001 *** | -1.34 | -5.4 |
|  | Inter<sub>TC</sub> | A<sub>TC</sub> | -4.53 | 36 | < 0.001 *** | -0.75 | -1.9 |
|  | AV<sub>TC</sub> | Inter<sub>TC</sub> | -5.23 | 36 | < 0.001 *** | -0.86 | -3.5 |
| Visual detriments | AV<sub>MC</sub> | A<sub>TC</sub> | 3.30 | 36 | 0.033 * | 0.54 | 1.5 |
|  | Inter<sub>MC</sub> | A<sub>TC</sub> | 1.01 | 36 | 1 | 0.17 | 0.4 |
|  | AV<sub>MC</sub> | Inter<sub>MC</sub> | 3.20 | 36 | 0.043 * | 0.53 | 1.1 |
| Coherence type benefits | AV<sub>TC</sub> | AV<sub>MC</sub> | -9.36 | 36 | < 0.001 *** | -1.54 | -6.9 |
|  | Inter<sub>TC</sub> | Inter<sub>MC</sub> | -5.54 | 36 | < 0.001 *** | -0.91 | -2.3 |
|  | A<sub>TC</sub> | A<sub>MC</sub> | -0.42 | 36 | 1 | -0.07 | -0.2 |
| Additional | AV<sub>TC</sub> | Inter<sub>MC</sub> | -8.75 | 36 | < 0.001 *** | -1.44 | -5.8 |
|  | AV<sub>TC</sub> | A<sub>MC</sub> | -9.18 | 36 | < 0.001 *** | -1.51 | -5.6 |
|  | AV<sub>MC</sub> | Inter<sub>TC</sub> | 6.66 | 36 | < 0.001 *** | 1.09 | 3.4 |
|  | AV<sub>MC</sub> | A<sub>MC</sub> | 3.34 | 36 | 0.029 * | 0.55 | 1.3 |
|  | Inter<sub>TC</sub> | A<sub>MC</sub> | -5.05 | 36 | < 0.001 *** | -0.83 | -2.1 |
|  | Inter<sub>MC</sub> | A<sub>MC</sub> | 0.68 | 36 | 1 | 0.11 | 0.2 |

### Video conditions can both enhance and impair participant performance

To more directly compare the influence of visual stimuli on listening, we calculated the visual benefits for each participant by comparing the AV_TC_ and Inter_TC_ with the A_TC_ condition (Figure 3B). Likewise, we visualised the effect of coherence on performance for each condition by comparing the thresholds for targets and maskers within each visual condition (Figure 3C).

The histogram representations of visual and coherence type benefits highlight these findings and offer insights into the variance of each benefit (Figure 3B, C). AV benefit values varied from −2 to 15.5 dB SNR, with 34 out of 37 listeners scoring a positive visual benefit for AV_TC_. The visual benefit in Inter_TC_ varied from −5.5 to 8 dB SNR, with 25 out of 37 participants achieving a positive value.

The impact of coherence on threshold also varied across condition (Figure 3C): Both AV and Inter coherence benefits were substantially positive, while the A coherence benefit was approximately symmetric around zero – consistent with the non-significant performance differences between A_TC_ and A_MC_ conditions. The AV coherence benefit varied from 0.5 to 15 dB SNR, the Inter coherence benefit from −2.5 to 8.5 dB SNR, and A coherence benefit from −3 to 6 dB SNR.

### Contributions of audiovisual temporal coherence and lipreading to the AV benefit are independent and vary across participants

To explore whether we could separate the contribution of lipreading to the AV benefit from other more general multisensory processes, we explored the relationship between AV and Inter benefits (Figure 4A). The correlation between the AV benefit and Inter benefit approached significance (Pearson’s r = 0.31, p = 0.058), but the borderline result is likely explained by the added variability introduced when computing benefit measures as differences between the original SRT_50_s.

**Figure 4:**
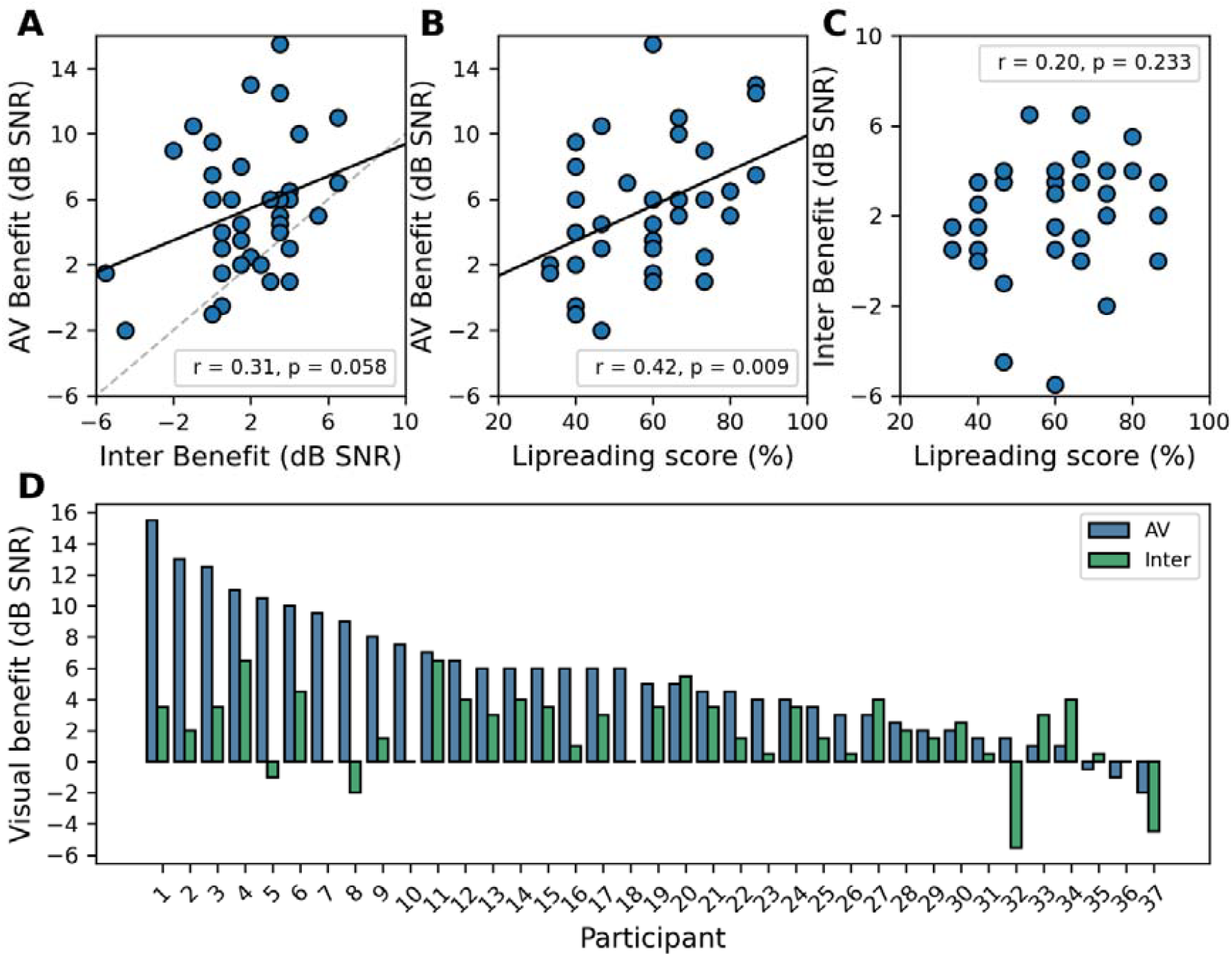
Linear dependencies and relative contributions of audiovisual temporal coherence and lipreading score to the AV benefit. A) Scatter plot and Pearson correlation between AV benefit and Inter benefit. B) Scatter plot and Pearson correlation between AV benefit and Lipreading score. C) Scatter plot and Pearson correlation between Inter benefit and Lipreading score. D) Bar plot displaying participant-specific AV (blue) and Inter (green) benefits, with participants ranked by the magnitude of their AV benefit. The height difference between the blue and green bars indicates the additional contribution of lipreading to the overall AV benefit. Black lines in A and B = regression lines; dashed line in A = equality line.

To further understand to what extent the AV benefit reflected lipreading ability, we asked to whether the independent TAS measure for sentences correlated with either the AV or Inter benefit. A Pearson correlation was conducted between AV benefit and Lipreading score (Figure B) revealing a small-to-moderate positive correlation (Pearson’s r = 0.42, p = 0.009). In contrast, the Inter benefit and Lipreading scores (Figure 4C) were not significantly associated (Pearson’s r = 0.20, p = 0.233), supporting the idea that the benefit in the Inter condition emerges from a separate multisensory process. To examine the relative contributions of both lipreading and non-lipreading mechanisms within individual participants, the AV and Inter benefits were plotted for each subject (Figure 4D; blue and green bars, respectively). Within this visualisation the lipreading contribution is reflected by the difference between the height of the AV benefit and Inter benefit bars. Figure 4D clearly shows substantial inter-individual variability in the relative use of the two mechanisms, including extreme cases where individuals obtain almost all of their AV benefit using only one of the two.

## Discussion

In the present study, we investigated how visual cues from a talker’s face affect speech-in-noise perception in young, normal-hearing participants. To do so, we developed a novel speech-in-noise test (vCCRMn) that incorporates both auditory and visual information. This task was specifically designed to capture the ‘AV benefit’ – that is, the performance improvement observed when individuals process audiovisual speech compared to audio-only speech – by quantifying contributions from both directly lipreading the target words and assisting with auditory selective attention more broadly.

Our results demonstrated that the vCCRMn reliably measured an AV benefit, as participants showed improved performance in the AV target-coherent (AV_TC_) condition compared to the audio with static image (A_TC_) condition. This finding aligns with previous studies reporting AV benefits for speech-in-noise perception (Bernstein & Grant, 2009; Grant & Bernstein, 2019; Sumby & Pollack, 1954).

Participants also performed significantly better in the Inter_TC_ condition compared to A_TC_. Because the Inter condition did not provide lipreading cues for the target words, the observed benefit cannot be attributed to lipreading of the target words and, instead, must relate to an enhanced ability for listeners to track the target talker through the mixture. What cues allow listeners to do this remains to be determined but is likely to be a mixture of correspondence between lip movements and speech, and low-level temporal coherence cues that promote audiovisual binding (Bizley et al., 2016; Lee et al., 2019; Maddox et al., 2015).

The benefit observed in the Inter_TC_ condition is consistent with visual information providing additional cues for speaker segregation beyond lipreading. Further evidence in favour of temporal coherence cues offering an additional benefit to that of lipreading was provided by the observation that while the AV benefit provided by the AV_TC_ condition was positively correlated with independently assessed lipreading scores, the Inter benefit did not.

Our scenes comprised of three co-located talkers, with one of the maskers being the same sex as the target talker, making segregation of the target talker from the mixture non-trivial. Audiovisual temporal coherence has been shown to confer a limited benefit when auditory scenes are simple to segregate (Cappelloni et al., 2023). As such, talker identity conveys a strong advantage on auditory selective attention performance, aiding, however, in selection rather than segmentation. In our study, we did not find significant differences between the two static image conditions (A_TC_ and A_MC_), ruling out the talker identity as a principal driver of the advantage seen in the Inter_TC_ condition. Another key difference between the study of Cappelloni et al. (2023) and those that have demonstrated advantages of audiovisual temporal coherence (e.g. Atilgan et al., 2018; Atilgan & Bizley, 2021; Maddox et al., 2015; Yuan et al., 2020, 2021) is that the latter studies all utilized amplitude cues to link audition and vision temporally, suggesting that this may be a key driver of both synthetic and naturalistic audiovisual binding.

In the Inter_TC_ condition of the vCCRMn, we chose to perturb lipreading by freezing the video during the colour and noun window of presentation, describing this manipulation to participants as ‘like a bad quality video call’. An alternative way to further separate articulatory correspondence and temporal coherence, utilized by Fiscella et al., (2022), could be to rotate the orientation of the face, thus potentially disrupting lipreading while maintaining temporal coherence. Such a control would be an interesting follow up to the work presented here, potentially revealing larger Inter benefits as temporal coherence cues might be better retained through the target word presentation period.

We also demonstrated that non-target visual information can impair speech-in-noise perception – performance in the AV_MC_ condition was significantly worse than in the audio-only condition. In the AV_MC_ condition, the audiovisual binding of the face and voice likely results in a stronger perceptual representation of the masker that both makes it harder to track the target talker through the mixture, and provides conflicting lipreading cues during the target presentation. The observation that there was no significant detriment elicited by the Inter_MC_ condition compared to audio only, and Inter_MC_ was significantly better than AV_MC_ suggests that lipreading during the target words is the key driver of this deficit, and in favour of the former interpretation.

As is common in studies of multisensory integration, there was considerable inter-individual variability in the magnitude of the AV benefit measured, and in the relative benefits observed for the full video (AV_TC_) and interrupted (Inter_TC_) conditions. Many participants derived some benefit in each condition, which is consistent with the interrupted condition capturing a subset of the available audiovisual benefit. However, some participants relied almost exclusively on Inter condition-derived benefits, while others not at all – with the latter, presumably, relying exclusively on lipreading-related cues for their AV benefits. This variability could have important clinical implications for populations who live with hearing loss, suggesting that personalised assessments of lipreading and audiovisual temporal coherence abilities may be valuable for designing effective rehabilitation and training programs.

## Acknowledgements

We thank Professor Mairéad MacSweeney for helpful discussions, Gordon Mills for implementing an online version of the vCCRM that allowed us to run pilot experiments during the COVID-19 pandemic, and the vCCRMn talkers for their help with recording the vCCRMn sentence stimuli. Finally, we thank the participants of the vCCRMn, without whom none of this work would have been possible.

## Declaration of Conflicting Interests

The authors declared no potential conflicts of interest with respect to the research, authorship, and/or publication of this article.

## Funding

This work was supported in whole, or in part, by the Wellcome Trust/Royal Society Sir Henry Dale Fellowship Grant 098418/Z/12/A and 227480/Z/23/Z to J.K.B.; the European Research Council Consolidator award 771550 SOUNDSCENE to J.K.B and a studentship from the NIHR UCL(H) Biomedical Research Centre (*Deafness and Hearing Problems* Theme) and the UCL Ear Institute to L.C.A.

## Notes

### Competing Interest Statement

The authors have declared no competing interest.

